# Transient and delay chemical master equations

**DOI:** 10.1101/2022.10.17.512599

**Authors:** Gennady Gorin, Shawn Yoshida, Lior Pachter

## Abstract

The serial nature of reactions involved in the RNA life-cycle motivates the incorporation of delays in models of transcriptional dynamics. The models couple a bursty or switching promoter to a fairly general set of Markovian or deterministically delayed monomolecular RNA interconversion reactions with no feedback. We provide numerical solutions for the RNA copy number distributions the models induce, and solve several systems with splicing and degradation. An analysis of single-cell and single-nucleus RNA sequencing data using these models reveals that the kinetics of nuclear export do not appear to require invocation of a non-Markovian waiting time.

## 1 INTRODUCTION

In spite of their substantial successes over the past few decades, Markov models may not be sufficient to describe biological processes. For example, consider the simplified RNA life-cycle depicted in Figure 1a: the unspliced RNA is transcribed from a DNA template, spliced within the nucleus, exported to the cytoplasm, and eventually degraded. There is considerable evidence that bacterial and mammalian transcription is typically Markovian and bursty (Dar et al., 2012, Fukaya et al., 2016, Lammers et al., 2020, Rodriguez and Larson, 2020, Sanchez and Golding, 2013), i.e., transcriptional events are apparently uniformly distributed in time, and generate multiple mRNA in rapid succession. This behavior is usually explained by appealing to a Markovian multi-state promoter model (Peccoud and Ycard, 1995); genes that exhibit more complex transcriptional dynamics are modeled by more sophisticated, but analogous Markovian models of transitions between promoter states (Ham et al., 2020b, Lammers et al., 2020, Zhou and Zhang, 2012). Similarly, the results of genome-wide inhibition experiments are largely consistent with Markovian RNA degradation, with exponentially decaying RNA levels over time (Sharova et al., 2009).

**Figure 1.**
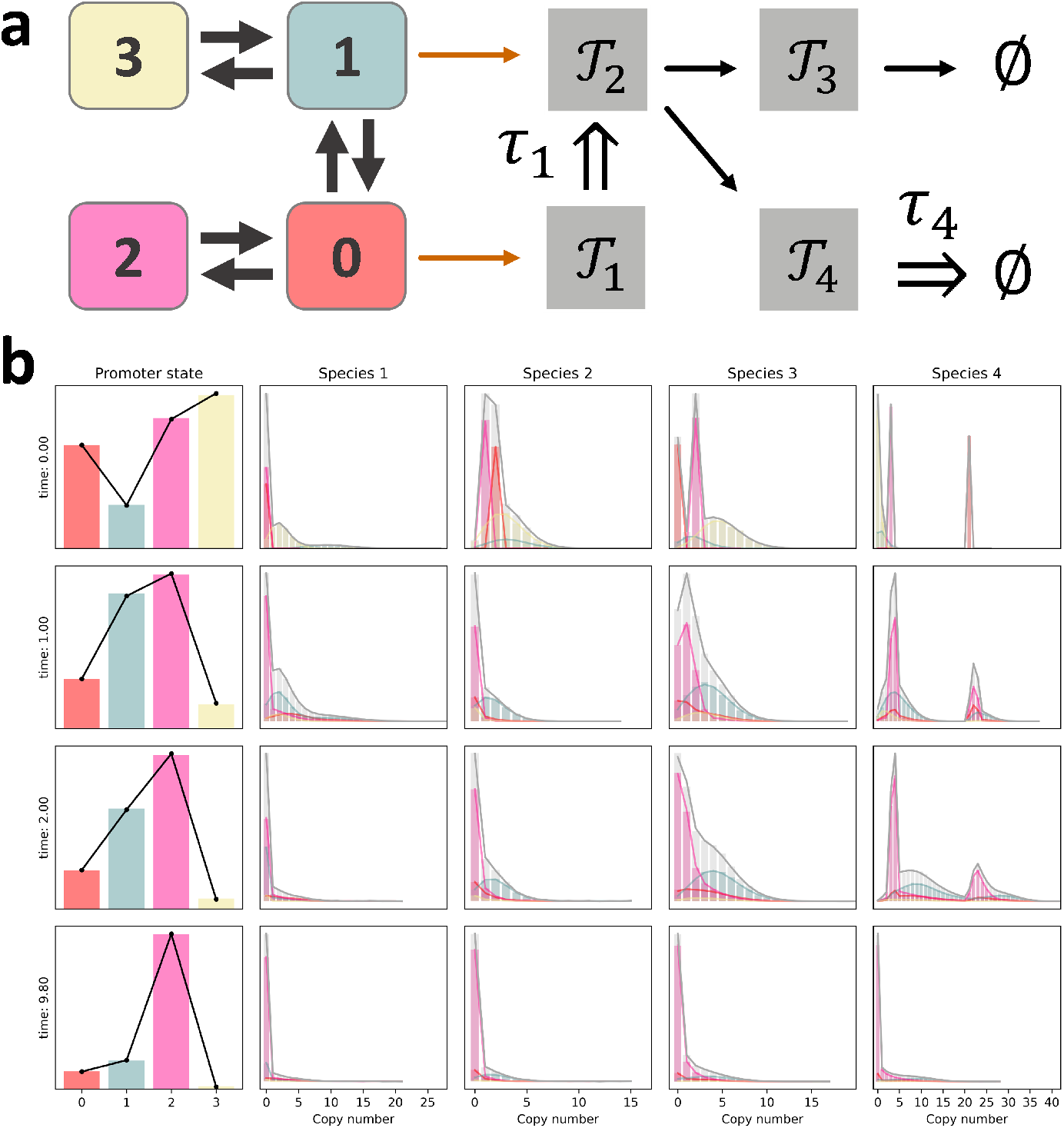
An outline of the experimental differences between single-cell and single-nucleus sequencing technologies. **a**. A schematic of our proposed model of the RNA life-cycle (gray outline: cell membrane; violet outline: nuclear membrane). **b**. The experimental workflows: single-cell sequencing uses whole cells, whereas single-nucleus sequencing uses isolated nuclei; both protocols produce a set of cDNA polymers that are sequenced and quantified to produce cell-by-gene matrices for the spliced and unspliced species. **c**. The differences in the experimental protocols translate to substantial and consistent differences in relative amounts of spliced and unspliced RNA: in single-cell datasets, unspliced RNA comprise 40–50% of the total dataset molecule counts, whereas in single-nucleus datasets, they predominate and comprise 70-90% of the total counts (colors: cell types; red: Allen mouse brain data; blue-green: Andrews human liver data). Illustrations adapted from (Gorin and Pachter, 2020b, Haque et al., 2017)

On the other hand, the kinetics of splicing and export are rather less well-characterized. For computational and mathematical tractability, our previous studies (Gorin and Pachter, 2021, 2022b) have elided the fine details of these processes. Thus, we typically assume that splicing is a single-step Markovian process, whereas export is rapid enough to neglect. These assumptions appear sufficient (Gorin and Pachter, 2021, 2022b) to fit data generated by single-cell RNA sequencing (scRNA-seq) that can quantify spliced (Zheng et al., 2017) and unspliced (La Manno et al., 2018, Melsted et al., 2021) RNA in whole cells (Figure 1b, gold and black arrows), but they have not been studied in detail.

The growth of single-cell RNA sequencing has been accompanied by advances in single-nucleus RNA sequencing (snRNA-seq) (Ding et al., 2020, Habib et al., 2017), which quantifies RNA in nuclei (Figure 1b, blue and black arrows). This technology does not provide a readout of cytoplasmic RNA, which occasionally reduces its utility for identifying genes of interest (Thrupp et al., 2020); however, it simplifies the experimental tissue handling, as scRNA-seq is incompatible with samples that are frozen or challenging to dissociate (Litviňuková et al., 2020, Selewa et al., 2020). The appropriate models for the genome-wide analysis of snRNA-seq data are not yet clear. We may plausibly retain the assumption that transcriptional and splicing dynamics are bursty and Markovian, consistently with single-cell data. However, we can no longer make the assumption of rapid export. As shown in Figure 1c, nuclear data are considerably enriched in unspliced RNA, but spliced RNA are present and the process of their export from the observed nuclear system must be mechanistically parameterized to fit the bivariate molecule distributions.

It is not *a priori* obvious that this export process should be Markovian. Previous theoretical studies have adopted this assumption (Filatova et al., 2022, Singh and Bokes, 2012), and some experimental studies involving synthetic fluorescent reporter constructs (Hansen et al., 2018, Munsky et al., 2018) have not observed gross discrepancies with the Markovian hypothesis. However, in previous work (Gorin and Pachter, 2022a), we observed that scRNA-seq data were largely consistent with the simple CME model of bursty transcription, Markovian splicing, and Markovian removal of spliced molecules, but snRNA-seq data exhibited a high rate of apparent model misspecification according to a goodness-of-fit criterion. We speculated that this phenomenon could be explained by a qualitative difference between mechanisms observable in single-nucleus and single-cell data. In single-cell systems, the spliced “efflux” step may reflect the relatively slow, Markovian degradation of spliced RNA, whereas in single-nucleus systems, it may reflect a more rapid, non-Markovian nuclear export process. However, the discrepancy may also be attributable to hard-to-model technical noise contributions (such as background counts introduced by the lysis process, or incompletely lysed cells erroneously sequenced along with nuclei), the violation of model assumptions (such as internal homogeneity), or the arbitrary choice of thresholds for the chi-squared test, necessitating dedicated model comparisons.

In the current study, we evaluate the consistency of the Markovian hypothesis in a more controlled fashion. We solve the models with deterministically delayed splicing and export, fit them for thousands of genes across five cell types from two species, and compare them to the standard Markovian model, using paired single-nucleus and single-cell datasets. We seek to answer the following three questions, ordered from least to most general:

- Do single-nucleus data *require* models that are substantially different from single-cell data, i.e., is the Markovian efflux hypothesis valid in nuclei?
- Given that transcriptional elongation is strongly non-Markovian (Corrigan et al., 2016), and splicing often (Coté et al., 2021) occurs co-transcriptionally (Drexler et al., 2020), is the Markovian splicing hypothesis consistent with sequencing data in general?
- To what extent do single-cell and single-nuclei data allow model identification?

Although the modeling, inference, and model discrimination procedures are quantitative, the answers we seek are largely qualitative. Furthermore, the strength of the conclusions is necessarily limited by the usual assumptions made for tractability, including the omission of technical noise, cell cycle, cell size effects, uncertainty in cell type assignments, and the granular structure of distinct intermediate unspliced transcripts. For simplicity, we do not attempt to treat the full scope of possible non-Markovian dynamics, and focus on deterministically-timed delays.

The schema in Figure 1a, as well as the considerations in the previous section, motivate a bursty promoter coupled to a two-stage model of RNA processing for nuclear and whole-cell data; as usual, we adopt the assumption of rapid efflux for scRNA-seq. To solve these models under the Markovian or non-Markovian hypotheses, we treat a considerably more general problem with an arbitrary number of downstream species and upstream promoter states, which potentially reduce to a set of co-bursting modules.

Extensive literature exists on the topic of stochastic delay systems. In particular, the stepwise elongation of a nascent mRNA strand during the process of transcription is frequently described as a deterministic delay (in the nomenclature of queuing theory, a *G / D / ∞* system) (Ali and Choubey, 2019, Choubey, 2018, Choubey et al., 2015, Fu et al., 2021, Xu et al., 2016, Zhang et al., 2014) or a series of small, stochastic steps along the template DNA strand (Gorin et al., 2020). Delayed regulation has been implicated in the p53 (Liu et al., 2011) and Hes1 systems (Barrio et al., 2006). Delay has been described as a regulatory motif in its own right, capable of modulating, suppressing, and buffering noise (Bhadana et al., 2021, Gedeon and Bokes, 2012, Singh et al., 2021, Zhang and Zhou, 2019), even beyond Markovian limits (Bhadana et al., 2021, Singh et al., 2021), as well as inducing oscillations (Bratsun et al., 2015, Liu et al., 2011, Miekisz et al., 2011, Singh et al., 2021).

In spite of these theoretical and biological applications, the studies described above generally discuss and derive statistical moments rather than full probability distributions, and few full analytical solutions are available for fitting transcriptomic copy number distributions. Several simple classes of delay chemical master equations (DCMEs) are solvable by the “linear chain trick” (LCT), which approximates a single constant delay of duration *τ* by *q* → ∞ exponentially-distributed delays with rate (*τq*) ^−1^ (MacDonald, 1978). At least in principle, the same idea can be used for any Erlang delay distribution (Jia and Kulkarni, 2011, Lafuerza and Toral, 2011b, Leier and Marquez-Lago, 2015, Zhang and Zhou, 2019). Unfortunately, the LCT is only practical when the distributions of all “virtual” intermediates are known in closed form, which is not the case for many biological systems of interest, such as bursty transcription (Gorin and Pachter, 2022a).

Several other analytical and numerical approaches have been proposed to solve DCMEs. There exists considerable literature on simulating delayed processes (Cai, 2007, Fatehi et al., 2020, Gupta et al., 2014, Tian et al., 2007), as well as approximating them by stochastic delay differential equations (SDDEs) (Gupta et al., 2014, Tian et al., 2007), which are typically more tractable. A recent report describes learning DCME solutions by designing a time-inhomogeneous CME with the time dependence of reaction rates encoded by a neural network (Jiang et al., 2021). Finally, reports by Lafuerza and Toral (Lafuerza and Toral, 2011a,b), Zhang and Zhou (Zhang and Zhou, 2019, Zhou and Zhang, 2012), Jiang et al. (Jiang et al., 2021), and Xu et al. (Xu et al., 2016) exploit the relationship between the DCME and closely related CMEs to obtain full probability distributions for several small systems.

In the current report, we find that this relationship is more generic and tractable than previously explored in the literature. We provide analytical and numerical recipes to determine the distributions of non-Markovian models with no feedback, and report and validate solutions for models with transcription governed by bursty and multi-state processes.

## 2 THEORETICAL FRAMEWORK

### 2.1 Preliminaries

We seek to model a system with *n* RNA species with counts 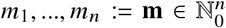. The molecules interconvert through monomolecular reactions, which are either Markovian, with waiting time distributed per Exp (*c*_*ik*_) for conversion from species *i* to species *k*, or delayed, with deterministic waiting time *τ*_*i*_. The molecules may also be degraded, with analogous allowed waiting time distributions. We assume the interconversion topology is described by a directed acyclic graph (Gorin and Pachter, 2022a).

We consider two models of transcriptional dynamics. In the first, we suppose the transcription process is entirely memoryless, and molecules are produced in bursts according to a Poisson arrival process with rate *α*_*i*_. This system is fully specified by a probability law *P* (**m**, *t*|*P*^0^, 0), with the initial condition defined by the distribution *P*^0^(**m**).

In the second, we propose the transcribing promoter exists in one of *N* states, with interconversion rates *H* _*jk*_ for transitions from state *j* to state *k*. Each state has a characteristic transcription rate *α* _*j*,*i*_, where *j* is the state index and *i* is the molecule index; transcription occurs according to a Poisson arrival process. This system is fully specified by a probability law **P** (**m**, *t*|**w**, **P**^0^, 0), where the entries of **P** contain the non-normalized probabilities *P* _*j*_ (**m**, *t*) for each promoter state. The initial conditions include **P**^0^, the set of non-normalized molecule distributions for each state imposed at *t* = 0.

Solutions for these systems typically cannot be obtained in closed form. However, they can be calculated by the Fourier inversion of the probability generating function (PGF), defined as follows:

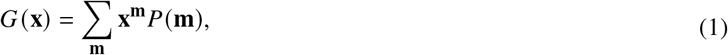

where the summation takes place over all microstates 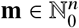, and **x^m^** is an informal shorthand for 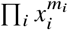. It is typically more straightforward to consider *G* (**u**), with *u*_*i*_ = *x*_*i*_ − 1. Further, the multi-state problem affords a PGF-like vector object **G**, with components *G_j_* such that

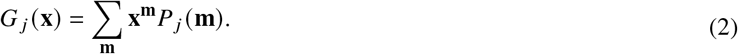

### 2.2 The transient dynamics of bursty systems

As previously discussed in (Gorin and Pachter, 2022a), *G*^*h*^, the PGF of a Markovian bursty process started at the homogeneous initial condition **m**_0_ = 0 has the following logarithm:

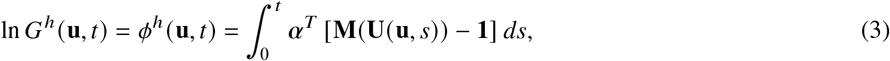

where the inner product is taken over all distinct combinations of burst processes in the system. The entries of ***α*** are the burst process arrival frequencies. The entries of **M** are the burst process moment-generating functions (MGFs). **u** is a vector of generating function arguments. The interconversion and degradation processes induce a series of *characteristics* **U** (**u**, *s*), which solve the following series of ordinary differential equations:

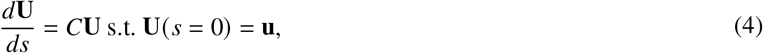

where 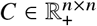 is a matrix containing the rates of interconversion and degradation of the *n* species.

To obtain the PGF of a non-homogeneous system, started at a distribution with the arbitrary PGF *G*^0^ (**u**), we take advantage of the fact that the newly transcribed molecules are statistically independent from the pre-existing molecules:

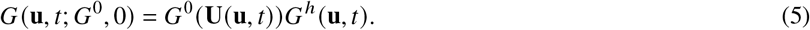

This formula can be evaluated using a single integral per value of **u**.

### 2.3 The transient dynamics of multi-state systems

It is straightforward to extend the foundations in (Gorin and Pachter, 2022a) to include interconversion between promoter states. We assume there are *n* species, with microstates 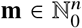, and *N* promoter states, with microstates *j* ∈ {0, …, *N* − 1}. The species are one-indexed; a reaction from a species to “0” corresponds to degradation. The promoter states are zero-indexed, to reflect the common notation wherein a “0” state is inactive whereas the “1” state is active.

We assume the system contains the following Markovian reactions:

- Interconversion between promoter states.
- Synthesis of new molecules.
- Interconversion of molecular species.
- Degradation of molecular species.

Define the collection of state-specific generating functions **G** ≔ [*G*_0_, …, *G* _*N*−1_]^*T*^, such that

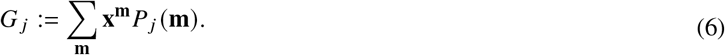

This collection is associated with the *N* × *n* Jacobian matrix, such that

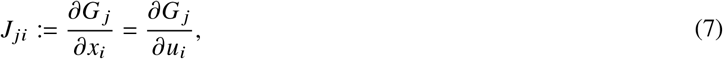

defining *u*_*i*_ = *x*_*i*_ − 1 for each generating function argument.

By definition, the master equation terms for switching between states contribute the following terms to the PDE system:

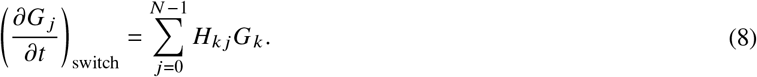

*H* ∈ ℝ^*N*^ is the matrix encoding the continuous-time Markov chain (CTMC) governing the promLoter states, such that Σ*_k_H_jk_* = 0 for all *j*. The diagonal terms encode the net efflux rate from state *j*, such that *H_jj_* ≔ Σ_*k*≠*j*_*H_jk_*. This set of reactions can be represented in the usual form for a finite CTMC:

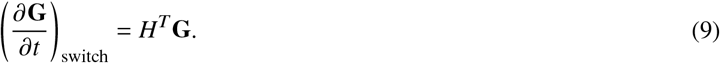

From (Gorin and Pachter, 2022a), the master equation terms for RNA synthesis contribute the following terms to the PDE system:

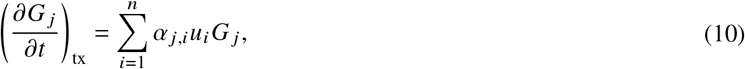

This set of reactions can be summarized by the matrix *A* ∈ (ℝ _≥0_)^*N*×*n*^, such that *A* _*ji*_ = *α _j,i_* is the rate of production of transcript *i* while the gene is in state *j* :

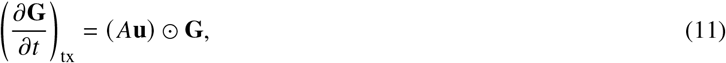

where the symbol ⊙ denotes the Hadamard, or entrywise, product of two matrices of identical dimensions.

Finally, the master equations for monomolecular, non-catalytic interconversion or degradation contribute the following terms to the PDE system:

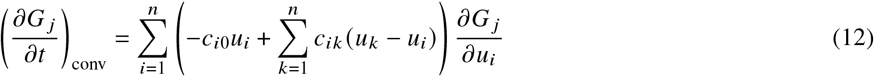

where *c_ik_* is the rate of transitioning from species *i* to species *k* and *c*_*i*0_ is the degradation rate of species *i*. This set of reactions can be summarized by the matrix *C* ∈ ℝ^*n*×*n*^ encoding the transitions between molecular species, such that *C*_*ik*_ = *c*_*ik*_ and *C_ii_* = −Σ_*k*≠*i*_*c_ik_* − *c*_*i*0_:

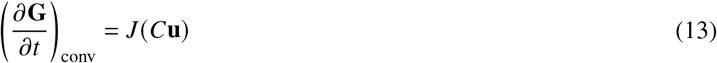

The coupled generating function PDEs take the following form:

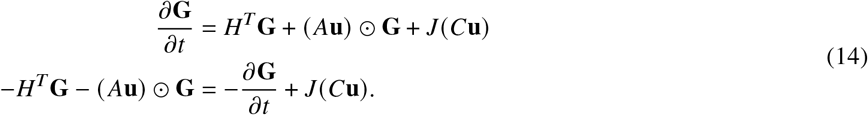

For each *j*, we can apply the method of characteristics (Singh and Bokes, 2012):

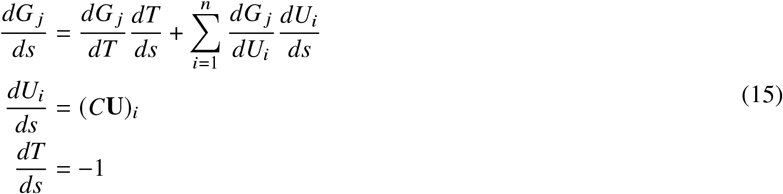

Imposing *T* (*s* = 0) = *t*, we find *T*(*s*) = *t − s*. By enforcing *U*_*i*_ (*s* = 0) = *u*_*i*_, we can simultaneously solve the system *∂*_*s*_**U** = *C***U**. We then yield:

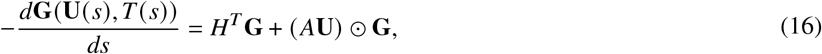

an initial value problem in *N* variables, parametrized by the characteristic variable *s*. We seek to obtain **G** at *T* = *t*, i.e., *s* = 0. We possess an initial condition **G**^0^ at *T* = 0, i.e., *s* = *t*. By numerically integrating −*H*^*T*^ **G** − (*A***U**) ⊙ **G** from *s* = *t* to *s* = 0, using **G**^0^ (**U**(*t*)) as the initial condition, we yield **G**(**u**, *t*).

#### 2.3.1 Solution through the lens of matrix ODEs

In practical terms, Equation 16 and its associated initial conditions are the solution to a particular switching system: the ODE can be easily plugged into a standard Runge–Kutta-type solver to obtain the generating function. However, we can, in principle, write down the ODE in more compact form:

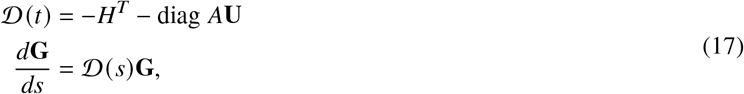

where 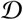 is the matrix that encodes system *d*ynamics. We observe:

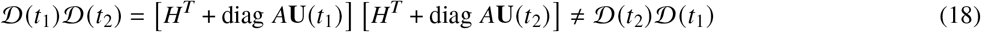

in general; the product of diagonal matrices commutes, but the product of *H*^*T*^ and a diagonal matrix does not (except in the trivial case *N* = 1). Therefore, **G** cannot be represented by a finite matrix exponential, but can be approximated by a Magnus series (Iserles and MacNamara, 2017).

#### 2.3.2 Solution through the lens of special functions

Certain steady-state solutions can be obtained by appealing to the theory of special functions. The stationary PGF of the solution to the standard telegraph model (with *N* = 2 and *n* = 1) is given by the confluent hypergeometric function _1_*F*_1_ (Iyer-Biswas et al., 2009); transient solutions have an analogous form. The combinatorial extension of the telegraph model (with *N* = 2^*p*^, *p* ∈ ℕ), whose state transitions are given by the Hamming graph, has a stationary PGF given by the *p*-fold product of confluent hypergeometric functions (Ham et al., 2020b). Certain systems with *n* = 1 and an irreversible refractory state dynamics have solutions given by the generalized hypergeometric function _*N* − 1_*F*_*N* − 1_ (Zhou and Zhang, 2012). Thus, in principle, solutions may be obtained by casting the system of *N* ODEs in Equation 16 into a single order-*N* ODE, then solving it using the properties of special functions. However, this tends to be considerably more involved than the original problem, both due to the complexity of manipulating hypergeometric functions and the challenges of actually evaluating them. Further, although hypergeometric representations are available for *n* = 1, the case of *n* > 1 does not even afford a formal solution. We have found the ODE approach yields a satisfactory combination of numerical stability and robustness for most purposes.

### 2.4 Applications to delay master equations

The numerical recipes outlined above can be exploited to solve delay chemical master equations. Instead of using Equation 4 to obtain the characteristics, we solve the downstream system by hand, using Equations S35 and S36 of (Gorin and Pachter, 2022a) for species with Markovian efflux and the identity

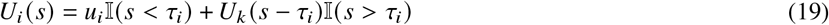

for species *i* converted to species *k* after a deterministic delay *τ*_*i*_. 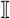 denotes the indicator function, which returns 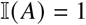 if *A* holds and 0 otherwise. If species *i* is degraded after a deterministic delay, its characteristic is simply 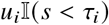. The resulting collection of characteristics can be plugged directly into Equation 5 or 16 and integrated.

## 3 RESULTS

### 3.1 Solving an arbitrary multi-state system

By numerical integration, we can obtain the molecule distributions induced by complex, arbitrary systems, such as the one shown in Figure 2a. To obtain the solution, we use Equation 16. We calculate **U**, use **G**^0^ (**U** (*t*)) as the initial condition to numerically compute **G**, and invert.

**Figure 2.**
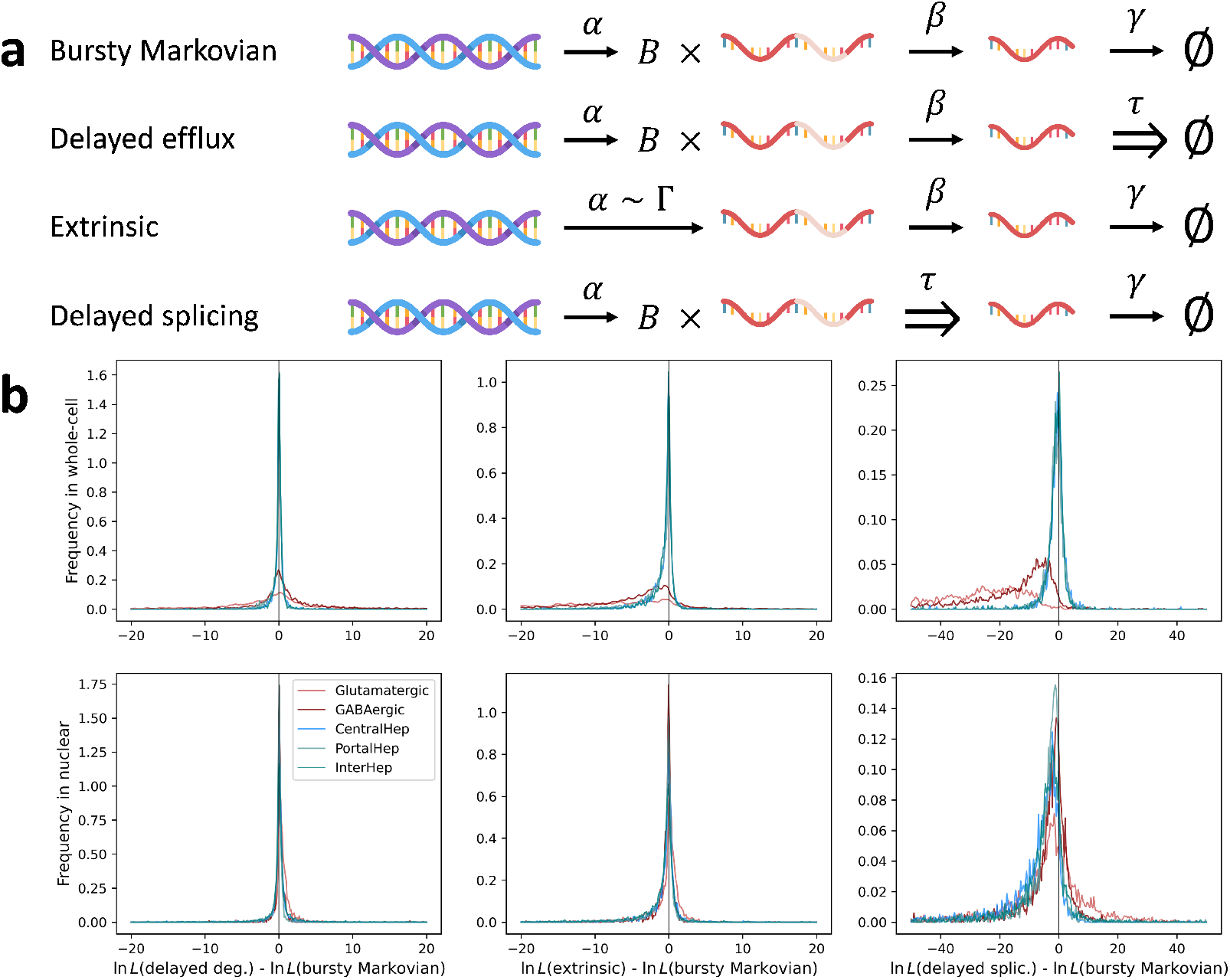
Validating a complicated non-Markovian transcriptional system. **a**. The structure of a system with four promoter states and four RNA species, two of which have deterministic lifetimes (rounded squares: promoter states; thick arrows: promoter state transitions; gray squares: RNA species; thin orange arrows: transcription reactions; thin black arrows: Markovian splicing and degradation reactions; wide arrows: deterministically delayed reactions). **b**. The analytical solutions for marginal distributions of promoter states and molecular species match simulations (Rows: time points. Leftmost column: promoter state distributions; histograms: simulation; lines: matrix exponential solution. Columns 2–5: the marginal distributions of molecular species; colored histograms: non-normalized marginals at each promoter state; gray histograms: molecular marginals aggregated over promoter states; lines: ODE solutions)

The parameters and initial conditions, given in Section 3.1.2, were set arbitrarily, taking care to keep the rates of interconversion reactions comparable and avoid regression to “trivial” regimes. The characteristics were computed using the MATLAB Symbolic Math Toolbox (MathWorks, 2022). As shown in Figure 2b, the procedure generates results that match simulations, and affords full conditional distributions at arbitrary times and for arbitrary initial conditions.

#### 3.1.1 Simulation and solution procedure

To compute the analytical solutions, we applied a standard Runge-Kutta integration method, implemented using `integrate.rk45` in the *SciPy* package (Virtanen et al., 2020).

To simulate the delayed system, we implemented a custom algorithm that introduces three modifications to Gillespie’s stochastic simulation algorithm (Gillespie, 1976). The Markovian form of the algorithm takes the following form:

1. Initialize system at time *t* = *t*_0_ and state *x* = *x*_0_.
2. Compute instantaneous reaction rates of the *μ*th reaction, flux_*μ*_ (*x*) and the net state efflux rate, flux(*x*) = Σ_*μ*_ flux_*μ*_ (*x*).
3. Compute sojourn time 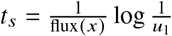 and reaction index, *μ*, such that 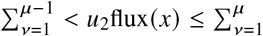, where *u*_1_ and *u*_2_ are random variables uniformly distributed over [0, 1].
4. Advance system to time *t ← t + t*_*s*_ and state *x ← x + ν*_*μ*_.
5. Return to step 2 or terminate simulation.

The first modification treats removal events of delayed species, i.e, transcripts that undergo reactions with deterministic waiting times. Two empty queues are initialized: one for times and one for reaction indices. Then, if the reaction index determined in step 3 is the creation of a delayed species, the queues are populated with the time and reaction index of the deterministic removal of that species in the correct order. If the system is initialized with delayed species, the queues of times and reaction indices must be pre-defined accordingly. For simplicity, we always assume that existing delayed species were created at *t* = 0.

The second modification alters the calculation of flux, specifically accounting for the contributions of species that don’t yet exist, but will after some delay. The total flux, flux *x*, is computed at each queued reaction event, and sojourn time corresponding to the random flux generated by 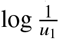 is found analytically. The computation of the sojourn time is essentially equivalent to the direct method outlined in (Cai, 2007).

The third modification ensures that all reactions happen in the correct order. After step 3, the reaction time and event are stored, and before advancing the system in step 4, all queued reactions that are to happen before the stored reaction event are sequentially applied and stored. The resulting ordered list is then converted into system times and states.

The algorithm is implemented in Python, and is designed to simulate the class of systems outlined in Section 2.3. With minor modifications, it can be used to simulate multi-molecular reactions, catalysis, feedback schema, or more general waiting time distributions, analogously to (Cai, 2007).

#### 3.1.2 Simulation parameters and characteristic solutions

The parameters for the system 2a are given in Table 1. The global parameters are *N* = 4 and *n* = 4. *H*_*i j*_ denotes the rate of transition from promoter state *i* to promoter state *j*. *α* _*j*,*i*_ denotes the rate of production of transcript 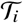 at promoter state *j*. *c*_*ik*_ denotes the rate of conversion of 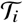 to 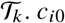 denotes the rate of degradation of 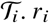 denotes the total efflux rate Σ_*k*_ *c*_*ik*_. *τ*_*i*_ denotes the residence time for deterministically delayed conversion. We generated realizations of trajectories for 10,000 cells, simulated until *t* = 10.

**Table 1.** Kinetic parameters used to simulate the multi-state DCME model

| Parameter | Value |
| --- | --- |
| $H_{01}$ | 0.75 |
| $H_{10}$ | 0.5 |
| $H_{02}$ | 1 |
| $H_{20}$ | 0.1 |
| $H_{13}$ | 0.15 |
| $H_{31}$ | 2 |
| $\alpha_{0,1}$ | 5.2 |
| $\alpha_{1,2}$ | 9.2 |
| $c_{23}$ | 1.8 |
| $c_{24}$ | 2.5 |
| $c_{30}$ | 0.8 |
| $\tau_1$ | 1.1 |
| $\tau_4$ | 2.1 |

The initial conditions are outlined in Table 2. The first row reports the probability of the system starting in each state. The following rows encode the distributions of each species at *t* = 0, conditional on starting in each state. D-*x* corresponds to a deterministic initial condition with *x* molecules, whereas P-*μ* corresponds to a Poisson initial condition with an average of *μ* molecules.

**Table 2.**
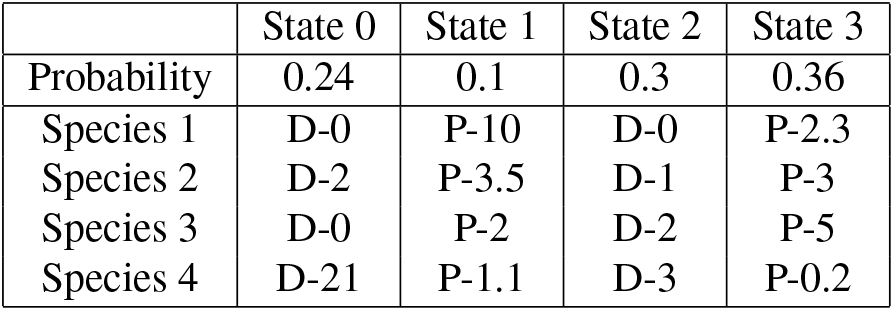
Initial conditions used to simulate the multi-state DCME model

|  | State 0 | State 1 | State 2 | State 3 |
| --- | --- | --- | --- | --- |
| Probability | 0.24 | 0.1 | 0.3 | 0.36 |
| Species 1 | D-0 | P-10 | D-0 | P-2.3 |
| Species 2 | D-2 | P-3.5 | D-1 | P-3 |
| Species 3 | D-0 | P-2 | D-2 | P-5 |
| Species 4 | D-21 | P-1.1 | D-3 | P-0.2 |

The system is described by the following characteristics:

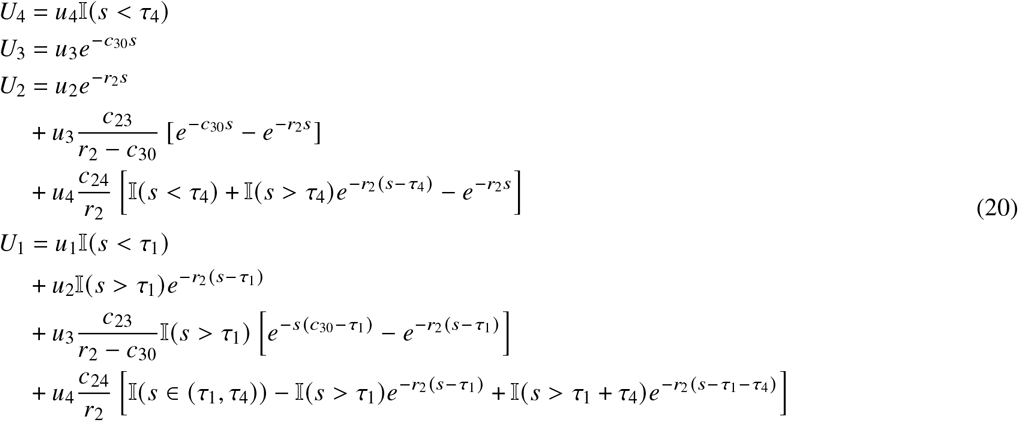

### 3.2 Single-nucleus data do not show identifiable mechanistic differences from single-cell data

As discussed in Section 1, we do not *a priori* know that splicing and nuclear export are Markovian. Using the suite of mathematical tools described in Section 2.2 and relatively homogeneous data, we revisit (Gorin and Pachter, 2021) the question of RNA life-cycle modeling and consider the differences between nuclear and whole-cell systems in further detail.

We are primarily interested in comparing the stationary distributions induced by the DCME reaction system with delayed efflux

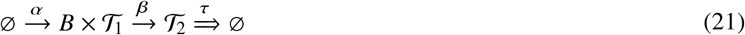

to those induced by the CME system

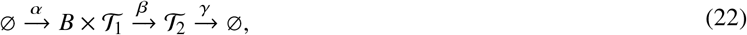

where *B* is distributed according to a geometric distribution on ℕ_0_ with mean *b*. We solve the DCME system in Section 3.2.2. In addition, we solve the closely related system with delayed splicing, or deterministic retention of the molecule at the gene locus, in Section 3.2.3:

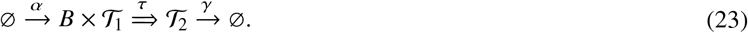

By using a computer algebra system, such as the MATLAB Symbolic Math Toolbox (MathWorks, 2022), it is straightforward to obtain the lower moments. For comparison, Table 3 displays the moments for the Markovian and delayed systems.

**Table 3.** Lower moments of the models of interest

| Moment | Markovian | Delayed efflux | Delayed splicing |
| --- | --- | --- | --- |
| $\mu_1$ | $\frac{\alpha b}{\beta}$ | $\frac{\alpha b}{\beta}$ | $\alpha b \tau$ |
| $\sigma_1^2$ | $\mu_1(1+b)$ | $\mu_1(1+b)$ | $\mu_1(1+2b)$ |
| $\mu_2$ | $\frac{\alpha b}{\gamma}$ | $\alpha b \tau$ | $\frac{\alpha b}{\gamma}$ |
| $\sigma_2^2$ | $\mu_2 \left( 1 + \frac{b\beta}{\beta+\gamma} \right)$ | $\mu_2 + 2\alpha b^2 \left[ \tau + \frac{1}{\beta} (e^{-\beta\tau} - 1) \right]$ | $\mu_2(1+b)$ |
| $\text{Cov}(X_1, X_2)$ | $\frac{\alpha b^2}{\beta+\gamma}$ | $\frac{\alpha b^2}{\beta} [1 - e^{-\beta\tau}]$ | 0 |

We fit the three models introduced above, as well as a qualitatively similar overdispersed model with a gamma-distributed transcription rate (Gorin and Pachter, 2020a, Gorin et al., 2021, Ham et al., 2020a) (Figure 3a), interpreting unspliced RNA counts as 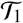 and spliced counts as 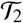. The analysis was performed using *Monod* (Gorin and Pachter, 2022b), using gradient descent to obtain maximum likelihood estimates (MLEs) under the four models (Section 3.2.4). As misspecification was evident with or without length bias in the original report (cf. `brain_5k_nuc` in Figs. S5 and S6 of (Gorin and Pachter, 2021)), we omitted the modeling of technical noise. The models were separately fit to GABAergic and glutamatergic cell types from two mouse brain samples (Yao et al., 2021), one generated using single-cell sequencing and one generated using single-nucleus sequencing, as well as single-cell and single-nucleus data from pericentral, periportal, and interzonal hepatocytes from a single human liver sample (Andrews et al., 2022).

**Figure 3.**
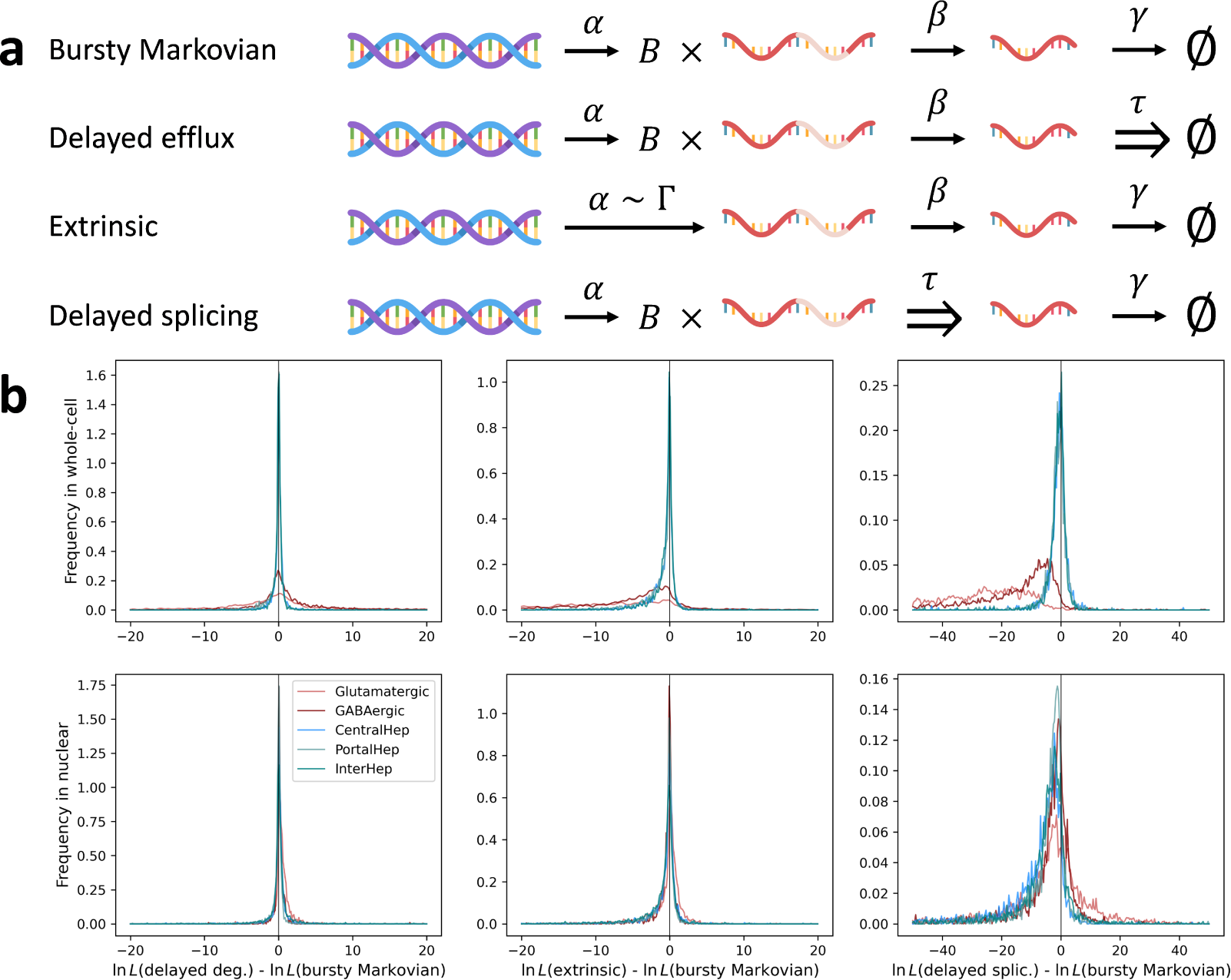
Certain models of RNA transcription and processing may be distinguished based on single-cell data, but single-nucleus data do not suggest that nuclear export biophysics are identifiably non-Markovian. **a**. The reaction schema of the considered models: DNA generates unspliced RNA with transcriptional burst frequency *α*, the unspliced RNA are converted to spliced RNA; the spliced RNA are removed from the system. In the “extrinsic” model, *α* is gamma-distributed; in the others, it is identical for all cells, and the transcriptional events produce bursts of size *B* ∼ Geom with mean *b*. **b**. Single-cell data (top row) support the bursty Markovian and delayed-efflux models to approximately the same degree, but offer consistently less support to the other two models, as quantified by the likelihood ratio. Single-nucleus data (bottom row) appear to show similar broad trends, but at lower magnitudes, limiting model identifiability (colors: cell types; red: Allen data; blue-green: Andrews data; lines: kernel density estimates)

Upon fitting the MLEs, we computed the models’ likelihood ratios relative to the standard bursty Markovian model and plotted them (Figure 3b). We did not observe a systematic bias toward the delayed-efflux model in the nuclear data (left column); the log-likelihood ratios were near zero in all considered cases, suggesting the data were insufficient to strongly favor either model in any of the datasets, nuclear or whole-cell. In contrast, the other two models were less favored relative to the standard model, with log-likelihood ratios tending to be negative. The strength of evidence against the models was lower in the single-nucleus data. This loss of statistical identifiability concords with intuition: the bursty Markovian, delayed-efflux, and extrinsic models afford identical negative binomial unspliced RNA marginals, requiring a large amount of spliced RNA to distinguish the models, which the nuclear sequencing protocol lacks. The delayed-splicing model has a different nascent marginal (described by the geometric-Poisson or Pólya–Aeppli distribution), and appeared to be more distinguishable from the negative binomial.

#### 3.2.1 Bursty Markovian system

The steady state of the system given in Equation 22 has been previously reported (Singh and Bokes, 2012) and has the log-PGF

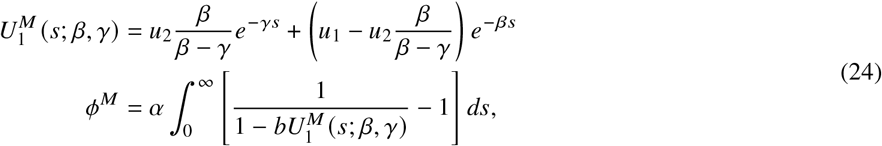

where 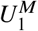 is the characteristic of the *M*arkovian system.

#### 3.2.2 Bursty with delayed efflux

The system given in Equation 21 is governed by the following equations:

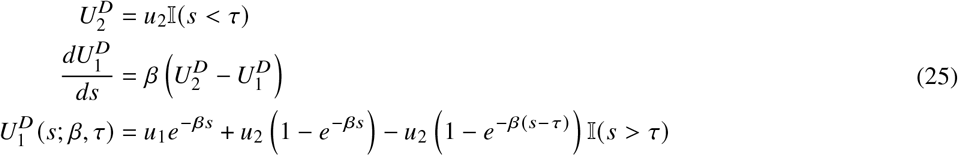

where 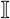 is the indicator function. This can be converted to a piecewise constant representation:

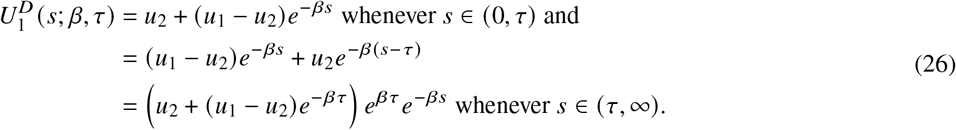

Defining *U* (*s*; *β*) ≔ *u*_2_ + (*u*_1_ − *u*_2_)*e*^−*βs*^ for convenience and integrating:

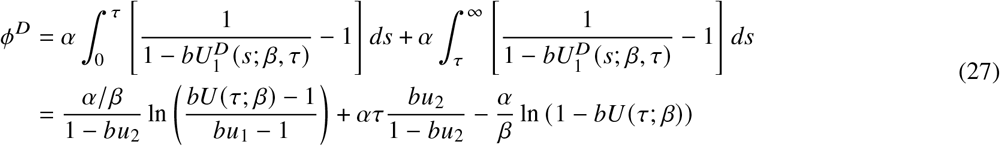

#### 3.2.3 Bursty with delayed splicing

The system given in Equation 23 supposes that each molecule of nascent RNA is retained in the system for duration *τ*. This is model is typical for studies concerned with modeling submolecular details of transcriptional elongation (Xu et al., 2016). This system is governed by the following characteristics:

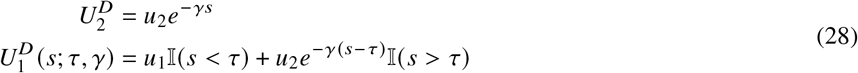

Upon plugging the characteristics into the burst size MGF and integrating, we find:

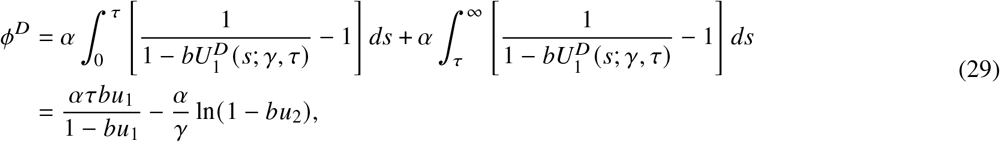

i.e., the joint distribution is merely the product of two independent marginal distributions. This result can be derived by appealing to the bursty limit of the argument in Section 14.4 in the supplementary material of (Xu et al., 2016).

#### 3.2.4 Data analysis

To compare the biophysical mechanisms consistent with single-cell and single-nucleus datasets, we used mouse cortex data generated by Allen Institute for Brain Science (“Allen data”) (Yao et al., 2021) and human liver data generated by Toronto’s University Health Network (“Andrews data”) (Andrews et al., 2022). The datasets analyzed for this study are reported in Table 4.

**Table 4.**
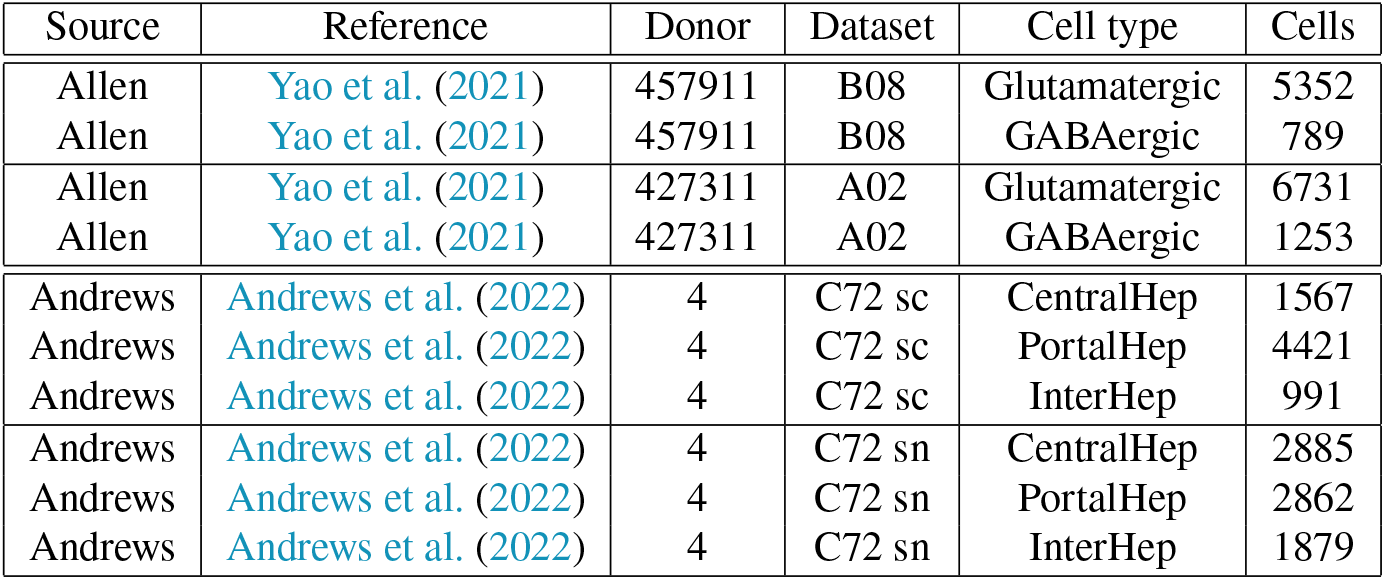
Sources and cell counts of analyzed cell type datasets

The Allen data did not contain technical replicates, i.e., a single mouse tissue processed using two single-cell and single-nucleus technologies. In lieu of this, we chose two libraries from female mice: B08, obtained from donor 457911, and A02, obtained from donor 427311. The raw single-cell FASTQs were obtained from https://data.nemoarchive.org/biccn/grant/u19_zeng/zeng/transcriptome/scell/10x_v3/mouse/raw/MOp/; the annotations were obtained from https://data.nemoarchive.org/biccn/grant/u19_zeng/zeng/transcriptome/scell/10x_v3/mouse/processed/analysis/10X_cells_v3_AIBS/. The raw single-nucleus FASTQs were obtained from https://data.nemoarchive.org/biccn/grant/u19_zeng/zeng/transcriptome/sncell/10x_v3/mouse/raw/MOp/; the annotations were obtained from https://data.nemoarchive.org/biccn/grant/u19_zeng/zeng/transcriptome/sncell/10x_v3/mouse/processed/analysis/10X_nuclei_v3_AIBS/.

For quantification, we used a pre-built mm10 reference genome released by 10x Genomics, obtained from https://support.10xgenomics.com/single-cell-gene-expression/software/downloads/latest, to generate cDNA and intron references using *kallisto|bustools* 0.26.0 (`kb ref` with option `--lamanno`). Both libraries were produced using the 10x Genomics v3 chemistry. We pseudoaligned the B08 and B02 libraries using the 10x v3 cell barcode whitelist and the default *kallisto|bustools* filter (kb count with options `--lamanno, −x 10xv3`, and `--filter bustools`) to generate spliced and unspliced count matrices. The data were further filtered, excluding all cells with too few total (spliced + unspliced) UMIs, with a threshold of 10^4^ for B08 and 8 10^3^ for A02. These thresholds were chosen manually based on knee plots.

To choose cells for analysis by *Monod*, we identified cell barcodes which were (1) assigned to either the GABAergic or glutamatergic cell type according to annotations, (2) passed the *kallisto|bustools* filter and knee plot filters, and (3) did not belong to one of the excluded rare cell subtypes (Scng or L6 IT Car3). This procedure yielded the cell numbers reported in the final column of Table 4.

The Andrews data contained technical replicates, generated using different sequencing chemistries. We analyzed data for donor 4 (`C72_reseq` for single-cell, here referred to as “C72 sc”; `C72_TST` for single-nucleus, here referred to as “C72 sn”). The single-cell data were obtained in the Sequence Read Archive format (experiment SRX12509405, runs SRR16227578–83) and converted to paired-end FASTQ files with *fasterq-dump* 2.11.2 (with options `--include-technical` and −S). The single-nucleus data were obtained in the originally submitted BAM format (experiment SRX12509406, run SRR16227584) and converted to paired-end FASTQ files with the 10x Genomics software *bamtofastq* 1.3.1. The metadata were obtained from the Gene Expression Omnibus (series GSE185477, file `GSE185477_Final_Metadata.txt.gz`).

For quantification, we used a pre-built GRCh38 reference genome released by 10x Genomics, obtained from https://support.10xgenomics.com/single-cell-gene-expression/software/downloads/latest, to generate cDNA and intron references using *kallisto|bustools* 0.26.0 (`kb ref` with option `--lamanno`). The single-cell library was produced using the 10x v2 chemistry, whereas the single-nucleus library was produced using the v3 chemistry. To simplify the procedure of extracting and storing the count data, we used the lists of barcodes present in each sample’s cell type annotations as the whitelists for pseudoalignment. The *kallisto|bustools* was run using the `--lamanno` option and the appropriate −x arguments for each dataset. Due to the whitelist definition, we did not use the default filter. Instead, we excluded all cells with too few total UMIs, with a threshold of 8 × 10^2^ for single-cell and 2 × 10^3^ for single-nucleus, and too many total UMIs, with corresponding thresholds of 10^4^ and 5 × 10^4^. These thresholds were chosen manually based on knee plots.

To choose cells for analysis by *Monod*, we identified cell barcodes which were (1) assigned to pericentral (CentralHep), periportal (PortalHep), or interzonal (InterHep) hepatocyte cell types according to annotations and (2) passed the knee plot filters. This procedure yielded the cell numbers reported in the final column of Table 4.

The analysis was performed using *Monod* 0.2.4.0. For each species, we selected 3,000 genes with moderate to high expression in all datasets. This procedure imposed a set of constraints (each species’ mean copy number > 0.01, each species’ maximum copy number > 3 and < 400) and computed the number of datasets in which the criterion was met. The top 3,000 genes were identified by choosing the top quantiles of this metric, using random non-replacement sampling to break ties.

Next, we fit all of the models to the data. The procedure used gradient descent to minimize the Kullback-Leibler divergence (KLD) between model and data, and output the maximum likelihood estimates (MLEs) for the parameters. To compute the proposed probability distributions for the KLD, we evaluated the generating functions on a two-dimensional grid of microstates, with the lower bound set to zero and the upper bound set at 10 molecules more than the observed maximum of each species. The generating function of the bursty Markovian model, which requires numerical integration, was evaluated by order-60 Gaussian quadrature, implemented through `integrate.fixed_quad` in *SciPy* (Virtanen et al., 2020), evaluated on the interval [0,*T*], where *T* = 10 (1 + β^−1^ + γ^−1^) to ensure the system reached equilibrium. For the bursty models, α was set to unity with no loss of generality at steady state. For the extrinsic noise model, *θ*, the scale parameter of the gamma distribution, was set to unity.

Gradient descent was performed using L-BFGS-B, a constrained optimization algorithm, implemented through the *SciPy* function `optimize.minimize` (Virtanen et al., 2020). The optimization was performed in a 3-dimensional space of log_10_ parameter values, with domain of 2, 3.1 for the burst sizes and gamma shape parameters and 1.8, 3.5 for *β*, *γ*, and *τ*^−1^. These parameters were chosen based on the parameter distributions seen in Figure S5 of (Gorin and Pachter, 2021). Optimization ran for a maximum of 15 steps, with 5 independent trials. The first trial was initialized at the method of moments (MoM) estimate. The method of moments estimates for the bursty Markovian models and the extrinsic models are reported in the supplement of (Gorin and Pachter, 2022b). The MoM estimate for the delayed efflux model was identical to that of the bursty Markovian model, with *τ*^−1^ taking the place of *γ*. The MoM estimate for the delayed splicing model was identical to that of the bursty Markovian model, with *b* corrected by a factor of 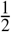 and *τ*^−1^ taking the place of *β*. These estimates follow directly from the results in Table 3.

The log-likelihood ratios visualized in Figure 3 were evaluated by computing the proposal PMFs as above, then computing the difference between the log-likelihoods of the data under the alternative model and under the bursty model:

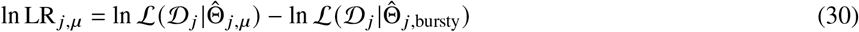

where *μ* is the alternative model type (delayed-efflux, extrinsic, delayed-splicing), *j* is the gene index, 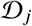 is the dataset (spliced and unspliced copy numbers in a particular cell type) for gene *j*, 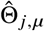 is the vector of parameter MLEs under the model, and 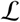 is the data likelihood.

Log-likelihood ratios with magnitude > 100 for single-cell and > 50 for single-nucleus data were omitted as potential signatures of failure to converge to a satisfactory MLE. Kernel density estimates in Figure 3b were generated from the remaining ln LR _*j*_ values using the SciPy class `stats.gaussian_kde`, with bandwidth set to 0.01 to avoid oversmoothing.

## 4 METHODOLOGICAL EXTENSIONS

In the current section, we draw useful connections to theory we previously reported in (Gorin and Pachter, 2021), contextualize the work with respect to standard tools of DCME analysis, and discuss potential extensions.

### 4.1 Generating function identifiability

Consider a system with *n* species, described by a generating function *G* (**u**). Some of these species may not be mutually identifiable. Suppose there are 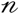 identifiable species, where each species *i* ∈ {1, …, *n*} may be assigned to each category 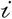 with probability 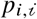 for each molecule. If this assignment process is independent, the process induces a collection of *n* generating functions 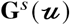, such that each component is the PGF of a categorical distribution (i.e., a multinomial distribution with a single trial):

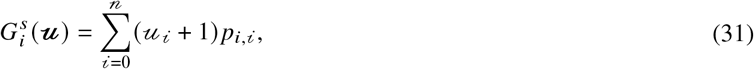

where *p*_*i*,0_ is defined as 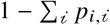, the probability of assigning species *i* to none of the observable species, and losing it. Evidently,

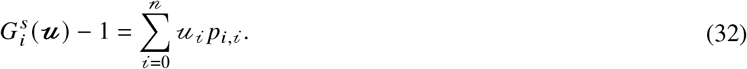

From standard properties of generating functions, the distribution resulting from “filtering” *G* through **G**^*s*^ is a function composition:

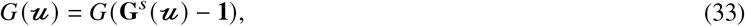

where **1** denotes the length-*n* vector of ones.

### 4.2 Erlang-distributed delays

Consider the case of a system with bursty transcription, Markovian degradation, and Erlang-distributed waiting time for splicing, with shape *q* and mean waiting time *τ*. First, set up a system with *q* + 1 species:

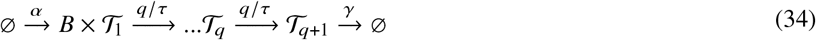

The characteristic corresponding to 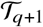 is simply *u*_*q*+1_*e*^−*γs*^. By somewhat tedious computation, amounting to repeatedly solving ordinary differential equations of the form 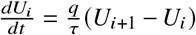, we find that the characteristic corresponding to 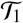 takes the following form:

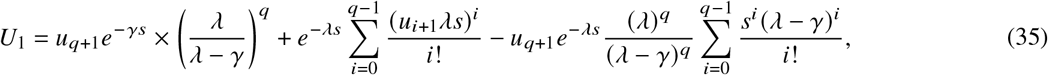

where *λ* is defined as *q /τ*.

To aggregate 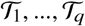 into a single species 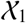, and represent 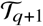 as the downstream species 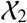, we evaluate the arguments at 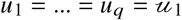 and 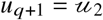. This yields:

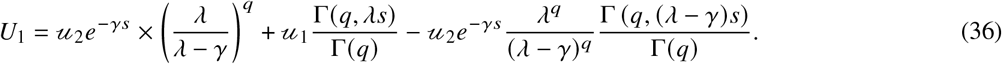

To evaluate the PGF, we plug *U*_1_ into the burst distribution MGF and integrate.

### 4.3 Linear chain trick

The same approach can be used to arrive at the solution for deterministically delayed systems, commonly known as the “linear chain trick.” Taking the limit as *q* → ∞ and exploiting the properties of special functions:

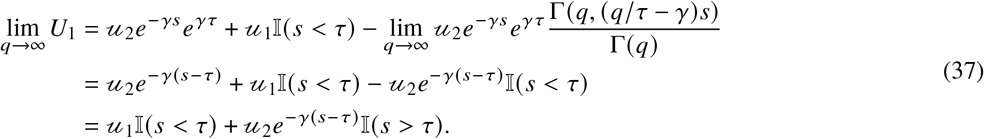

To obtain the PGF, we plug lim_*q*→∞_ *U*_1_ into the burst distribution MGF and integrate, eliding the functional analysis justification for interchanging the limit and the integral. This reproduces the results in Section 3.2.3, albeit with considerably more effort.

### 4.4 General waiting time distributions

The same approach may be used to evaluate systems with more generic waiting time distributions. For example, the characteristic appropriate for a molecule that remains in the system for time *τ*, then exhibits Markovian efflux is immediately implied by the results of the previous section:

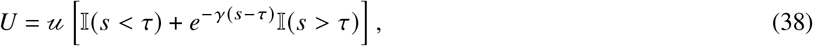

obtained by evaluating the previous characteristic *U*_1_ at 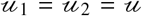. This holds for any combination of Markovian and deterministic delays.

Throughout the current section, we have considered delays only in downstream molecular processing. However, studies have indicated that non-Markovian refractory periods can occur in promoter state transitions (Harper et al., 2011). If the refractory period can be effectively described by a series of Markovian processes (e.g., if the waiting time is Erlang), such systems can be solved by defining “internal” inactive states and adding their *P* _*j*_, obtained through Equation 16, in the spirit of (Herbach, 2019). Stinchcombe et al. treated the case of arbitrary switching time distributions (Stinchcombe et al., 2012); however, this approach requires a degree of ingenuity and is fairly challenging to formalize in the language of generating functions.

The case of more general waiting times was explored in considerable detail by Zhang and Zhou (Zhang and Zhou, 2019), albeit with somewhat more emphasis on theoretical foundations and implications for noise buffering than here. Essentially, arbitrary waiting time distributions may be encoded by properly exploiting their statistical structure; for example, it is immediately evident that the characteristic in Equation 38 is simply the survival function (complementary cumulative distribution function) of each molecule, evaluated at its birth. In the same vein, the 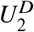 terms in Equations 25 and 28 are the survival functions of the degenerate and exponential distributions, whereas the *u*_2_ term of the characteristic in Equation 24 is the survival function of the two-parameter hypoexponential distribution.

It appears that more general waiting time distributions can be incorporated in the characteristic framework by appropriately manipulating their survival functions. We forgo detailed discussion of this extension, as it is less suitable to automation and tends to lead to intractable integrals. Further, Markovian and deterministic delays are relatively easy to motivate from first principles by appealing to memorylessness or perfect molecular memory, respectively (either maximum or zero entropy in the language of statistical physics (Pressé et al., 2013)). On the other hand, “intermediate” non-Markovian cases require a specific hypothesis to instantiate a functional form for the waiting time distribution.

The simulation algorithm outlined in Section 3.1.1 generalizes to arbitrary waiting time distributions, and may be used to investigate the impact of different assumptions even when analytical solutions are unavailable. Further, the tools outlined here can be used to explore some pathological cases, such as the loss of stationarity due to particularly ill-behaved waiting time distributions. For example, if the RNA production process is constitutive with unity transcription rate and the waiting time distribution is standard half-Cauchy, the characteristic and log-PGF take the following form:

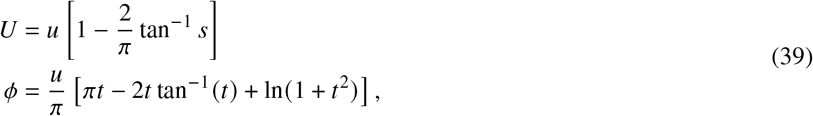

where *ϕ* is immediately recognizable as the log-PGF of a Poisson distribution, albeit one that does not possess a limit as *t* → ∞. This implies that the *M*/*G*/∞ queue with half-Cauchy service times has no stationary distribution. In contrast, if the waiting time distribution is standard Pareto, with support on [1, ∞), we find:

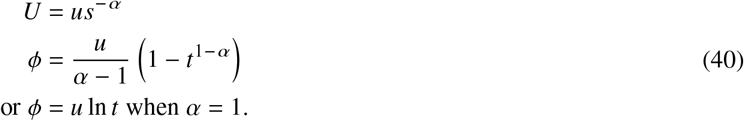

This distribution possesses a Poisson steady state with mean (*α* − 1) ^−1^ whenever *α* > 1. This implies that the *M / G /∞* queue with Pareto service times has a stationary distribution, but only if the service time distribution is endowed with a mean. Approaches such as this may be of use in identifying classes of non-Markovian processes admissible under particular axioms about biology, e.g., the existence of steady states, without assuming the existence of all Laplace transforms for the waiting time distributions (Zhang and Zhou, 2019).

## 5 DISCUSSION

This report extends the solutions reported in (Gorin and Pachter, 2022a) to a broad class of delayed systems, and unifies them with the theory of switching genes outlined in (Vastola, 2021). The approach augments the toolbox of stochastic biophysical analysis: it affords generating function-based analytical solutions for systems which would otherwise be outside the purview of the standard analysis of Markov chains. The derivation of these solutions does not require the usual manipulations of master equations at several time points, merely fairly standard integrals. These solutions may be fit (e.g., through the *Monod* implementation) and used as alternative hypotheses to investigate whether the standard assumptions of memorylessness hold, whether they are violated, or whether the available data are equivocal.

In Section 3.2, we used these analytical solutions to fit several superficially similar bivariate overdispersed models to single-cell and single-nucleus data, and observed that the typical Markovian model is a strong candidate, but cannot be easily distinguished from the competing delayed-efflux model, especially in the low-spliced expression single-nucleus data. On the other hand, two alternative models are consistently judged less plausible. This suggests that the assumption of Markovian, one-step splicing is useful despite its simplicity. This investigation is far from comprehensive. We forgo modeling technical noise or identifying cell types; using cell type definitions based on the spliced RNA counts introduces some degree of circularity. However, even with these limitations, the results illustrate fundamental challenges associated with single-nucleus data. To sum up, the data analysis is consistent with the following answers to the questions posed in Section 1:

- Single-nucleus data do not appear to require qualitatively different models. Both single-cell and single-nucleus data are consistent with the delayed efflux model.
- The Markovian splicing hypothesis is considerably more consistent with data than the delayed-splicing hypothesis.
- As may be expected, the low amounts of spliced RNA in nuclear data mean that only models with substantial differences in the unspliced distributions can be distinguished.

Furthermore, even when generating functions cannot be obtained, which is the typical case for systems with multiple RNA species, our method provides a simple numerical recipe for evaluating their distributions. The numerical approach is guaranteed to run in 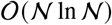, can use off-the-shelf integration routines (e.g., quadrature for bursty systems and Runge–Kutta methods for switching systems), and enables the evaluation of arbitrary marginal distributions, which is not feasible for matrix- or simulation-based methods.

The approach is largely modular with respect to the specific details of the downstream processes, i.e., the causal relationships between molecules and the Markovian or deterministic waiting times for the reactions. In Section 4, we discuss a set of extensions to even more generic systems, with waiting times described by combinations of exponential and degenerate distributions. The same generating function-based approach can be used to implement models of technical noise and molecular non-identifiability under the assumption of independent sampling (Gorin and Pachter, 2021). Although the solutions are fairly generic, they do not yet provide a route to treating more complex systems with protein translation or molecular feedback. We speculate that a sufficiently general treatment of such systems may uncover similar relationships between CMEs and DCMEs, and provide a simulation routine that can be extended to model such phenomena.

## Acknowledgments

G.G. thanks Dr. John J. Vastola and Catherine Felce for valuable discussions. G.G. and L.P. were partially funded by the National Institutes of Health grant U19MH114830. A part of the reported results were obtained during a Data Sciences Co-op with Celsius Therapeutics, Inc. The DNA and RNA illustrations are derived from the DNA Twemoji by Twitter, Inc., used under CC-BY 4.0.

